# Mapping numerical perception and operations in relation to functional and anatomical landmarks of human parietal cortex

**DOI:** 10.1101/602599

**Authors:** Elisa Castaldi, Alexandre Vignaud, Evelyn Eger

## Abstract

Human functional imaging has identified the middle part of the intraparietal sulcus (IPS) as an important brain substrate for different types of numerical tasks. This area is often equated with the macaque ventral intraparietal area (VIP) where neuronal selectivity for non-symbolic numbers is found. However, the low spatial resolution and whole-brain averaging analysis performed in most fMRI studies limit the extent to which an exact correspondence of activation in different numerical tasks with specific sub-regions of the IPS can be established. Here we disentangled the functional neuroanatomy of numerical perception and operations (comparison and calculation) by acquiring high-resolution 7T fMRI data in a group of human adults, and relating the activations in different numerical contrasts to anatomical and functional landmarks on the cortical surface. Our results reveal a functional heterogeneity within human intraparietal cortex where the visual field map representations in superior/medial parts of IPS and superior parietal gyrus are involved predominantly in numerosity perception, whereas numerical operations predominantly recruit lateral/inferior parts of IPS. Since calculation and comparison-related activity fell mainly outside the field map representations considered the functional equivalent of the monkey VIP/LIP complex, the areas most activated during such numerical operations in humans are likely different from VIP.

## Introduction

Learning mathematics is a hierarchical process in which every new concept builds on previously acquired knowledge and on lower-level cognitive primitives (Iuculano and Menon, 2018). One widely influential theory of numerical cognition proposed as the very first building block of mathematical knowledge the preverbal capacity to quickly determine the number of items in an image, based on the so called ‘number sense’ (Dehaene, 1997), that humans share with other animals since very early in life (Cantlon, 2012; de Hevia et al., 2017). During development, non-symbolic quantities are thought to be associated with symbolic numerals and the ability to fluently manipulate numerical magnitudes in these various formats, for example in the context of a numerical comparison task, has been shown to predict more complex arithmetical abilities (Anobile et al., 2016, 2018; Halberda et al., 2008; Libertus et al., 2011, 2013; Bugden et al., 2012; Holloway and Ansari, 2009; Sasanguie et al., 2012; Price and Fuchs, 2016; Chen and Li, 2014; Fazio et al., 2014; Schneider et al., 2017; Starr et al., 2017).

The predictive relationship at the behavioral level suggests the existence of a common neuronal substrate supporting numerical perception and different types of numerical operations. According to the triple code model the intraparietal cortex, and more specifically the horizontal segment of the intraparietal sulcus (HIPS), hosts a core module for processing the semantic meaning of numbers which is activated when perceiving or operating on numbers (Dehaene et al., 2003). This theory has received support from many fMRI studies showing that parietal areas, especially within and around HIPS, are recruited in a variety of diverse tasks involving numerical processing (Dehaene et al., 2003; Hubbard et al., 2005; Piazza and Eger, 2016; Eger, 2016; Knops, 2017). For example, areas within and around HIPS were found to be overall activated during approximate and exact calculation compared to non-numerical control tasks (Dehaene, 1999; Knops et al., 2009; Pinel and Dehaene, 2010; Pinel et al., 2007; Simon et al., 2002), as well as during numerical comparisons, where their BOLD signal was moreover modulated by the numerical distance of the compared numbers (Pinel et al., 2001, 2004; Ansari et al., 2006a). However, independent of the execution of such numerical operations, enhanced activity for numbers as opposed to letters or colors was also to a lesser extent measured in HIPS during an orthogonal target detection task (Eger et al., 2003). Moreover, parietal regions were reported to habituate to repeated presentation of the same numerical quantity and show numerical distance-dependent recovery of activity for deviant numbers (Piazza et al., 2004; Cantlon et al., 2006; Ansari et al., 2006b) to some extent even across formats (Piazza et al., 2007; Vogel et al., 2017), to encode numerical quantity in multi-voxel patterns of evoked activity (Borghesani et al., 2018; Bulthé et al., 2014; Castaldi et al., 2016, 2019; Cavdaroglu and Knops, 2018; Damarla and Just, 2013; Eger et al., 2009, 2015; Lasne et al., 2018) and to contain topographically organized numerosity maps (Harvey et al., 2013; Harvey and Dumoulin, 2017a, 2017b). Recently, two meta-analyses quantified the degree of overlap of the parietal activations elicited by different numerical tasks and concluded that the very same regions are recruited, namely the inferior and superior parietal lobules (IPL and SPL) which delimit the intraparietal sulcus (IPS), during calculation and non arithmetics-related numerical task, both in adults (Arsalidou and Taylor, 2011) and in children (Arsalidou et al., 2018).

FMRI studies in developmental dyscalculia showed abnormal activations of, among others, the mid-posterior parietal cortex during symbolic (Mussolin et al., 2010) and non-symbolic (Bulthe et al., 2018; Kaufmann et al., 2009a; Price et al., 2007) numerical comparisons, ordinality judgements (Kaufmann et al., 2009b), approximate calculation (Kucian et al., 2006) or simple arithmetical verification (Ashkenazi et al., 2009; Iuculano et al., 2015; Rosenberg-Lee et al., 2015) in dyscalculic children with respect to controls. Moreover, a meta-analysis identified the IPS as one of the areas consistently differing between individuals with and without dyscalculia during diverse number processing tasks (Kaufmann et al., 2011). Thus, overall, at least when considered at a coarse spatial scale, a large body of imaging work in humans suggests the existence of a neuronal substrate supporting the processing of number in human intraparietal cortex that is activated across a wide range of numerical tasks, and altered in subjects with impaired numerical skills.

Of note, similarly located regions, in the fundus of the intraparietal sulcus, were also found to be involved in numerical processing in non-human primates. More specifically, electrophysiological studies have recorded numerical responses of single neurons, which distinguish between different numbers of items presented, from ventral (VIP) and lateral (LIP) intraparietal areas in macaques (Nieder et al., 2006; Roitman et al., 2007). One influential review article compared the functional organization of human and macaque intraparietal cortex with respect to numerical processing and other response properties (Hubbard et al., 2005). These authors noted that in humans numerical processing-related activation foci (for estimation, comparison and simple arithmetic) were found in close spatial proximity to activations elicited by visuo-tactile multisensory, grasping and saccadic eye movement tasks, tasks that in monkeys activate areas VIP, AIP and LIP (Bremmer et al., 2001; Sereno et al., 2001; Simon et al., 2002). Based on these colocalizations and overall similarities with the spatial arrangement of the intraparietal sub-regions in macaques, a tentative homology was proposed between the brain regions activated for numerical tasks in humans, and macaque VIP (and to a lesser extent, LIP). However, beyond the uncertainties inherent in comparing brain locations in normalized space across groups of human subjects from different studies, and even across species, it is important to note that the numerical responses considered here in humans (mostly including the execution of numerical operations) were quite different from the ones investigated by macaque neurophysiology (which correspond to preferential responsiveness to different non-symbolic numbers). Therefore, it still remains to be confirmed whether at a more fine-grained level of anatomical localization of activations, these different aspects of numerical processing recruit identical sub-regions even in the same species, and what is their precise relation to the known functionally defined areas.

Arguably most similar to the non-symbolic number selectivity observed in macaque monkeys are the numerosity maps more recently discovered in humans, where individual voxels respond preferentially to different numbers of visual items (Harvey et al., 2013). In another recent review article, Harvey et al (2017) noted that in humans these maps were located more superior/medially in the superior parietal lobule, rather than in the fundus of the IPS where activations elicited by numerical comparison and calculation tasks usually appear to be centered. Based on these observations, they proposed that the neuronal circuits supporting basic physical quantity processing, i.e., numerosity perception, and numerical operations, as for example comparison, may be distinct, and questioned the often assumed functional homology between a core number processing system, localized in the human HIPS, and macaque VIP. However, this proposal was based on a review of the local maxima reported across multiple studies in different groups of subjects, where data were in addition acquired at different spatial resolutions and field strengths. Group analyses in whole brain space usually apply a considerable amount of volume-based smoothing and are likely to insufficiently represent the precise cortical location of activation foci. In addition, whole brain group maxima depend on the inter-subject variability of every given sample of subjects from which they are derived, and projecting such maxima from different studies onto an average surface could yield different locations due to the variability between the samples used, which need not reflect true differences in activated anatomical location.

In the work reported here, we explicitly tested, within the same group of human subjects, the idea that there exists a regional specialization within human intraparietal cortex with separate subregions recruited during different aspects of numerical processing including numerosity perception on the one hand and different numerical operations (comparison and mental calculation) on the other hand. For a more precise anatomical localization of activations, we exploited the enhanced resolution of ultra-high field (7T) fMRI in combination with extraction of the cortical surface in each subject. We further related the observed activations on the cortical surface to anatomical and functional markers derived from two atlases: one based on identifying the major sulci and gyri (Destrieux et al., 2010), and the other based on visual topography (Wang et al., 2015). The latter defines six areas, labelled from IPS0 to IPS5, which have been identified by means of phase-encoded mapping in human intraparietal cortex, from its most posterior to most anterior subparts (Konen and Kastner, 2008; Sereno et al., 2001; Silver et al., 2005; Swisher et al., 2007). Based on similarities in the relative anatomical localization and functional response properties, tentative homologies have been proposed between some of these field maps and the macaque LIP/VIP complex (Kastner et al., 2017; Konen and Kastner, 2008). If in humans there should be a single core system recruited to support both numerical perception and numerical operations, which is equivalent to the monkey LIP/VIP complex, we would expect activations to be centered within the visual field map representations in all conditions. On the contrary, if numerosity perception, numerical comparison and calculation rely on separable neuronal substrates within intraparietal cortex, our methods should allow us to better localize the sub-regions recruited by these different aspects with respect to the anatomy of intraparietal cortex, and refine the current ideas about which of these types of processing can be considered to occur in functionally homologous areas across species.

## Methods

### Subjects, data acquisition procedure and fMRI paradigms

Sixteen healthy adult volunteers (six males and nine females, mean age 25±2 years) with normal or corrected vision participated in the study. The experiment was approved by the regional ethical committee (Hôpital de Bicêtre, France) and undertaken with the understanding and written consent of each subject.

A SIEMENS MAGNETOM 7T scanner with head gradient insert (Gmax 80mT/m and slew rate 333T/m/s) and adapted 32-channel head coil (Nova Medical, Wilmington, MA, USA) was used to collect functional images as T2*-weighted Fat-Saturation echo-planar image (EPI) volumes with 1.3 mm isotropic voxels using a multi-band sequence (Moeller et al., 2010) (https://www.cmrr.umn.edu/multiband/, multi-band [MB] = 2, GRAPPA acceleration with [IPAT] = 2, partial Fourier [PF] = 6/8, matrix = 150 × 150, repetition time [TR] = 1.75 s, echo time [TE] = 21 ms, echo spacing [ES] = 0.74 ms, flip angle [FA] = 65°, bandwidth [BW] = 1516 Hz/px, phase-encode direction anterior to posterior). Calibration preparation was done using Gradient Recalled Echo (GRE) data. Fifty transversal slices covering the parietal and frontal cortex were obtained in ascending interleaved order. At the beginning of the scanning session, two single volumes were acquired with the parameters listed above but with opposite phase encode directions. The single-band reference images of these two initial volumes were used for distortion correction (see Data Analysis).

Anatomical images (T1-weighted) were acquired at 1 mm isotropic resolution using an MP2RAGE sequence (GRAPPA acceleration with [IPAT] = 3, partial Fourier [PF] = 6/8, matrix = 256 × 256, repetition time [TR] = 5 s, echo time [TE] = 2.82 ms, time of inversion [TI] 1/2= 800/2700 ms, flip angle [FA] 1/2 = 4°/5°, bandwidth [BW] = 240 Hz/px). A radiofrequency absorbent jacket (Accusorb MRI, MWT Materials Inc., Passaic, NJ, USA) was used to minimize the so-called “third-arm” or “shoulder” artifacts due to regions where the head gradient is unable to unambiguously spatially encode the image (Wald et al., 2005). The participants’ head was stabilized by padding and tape to prevent excessive movements. They saw the visual stimuli back-projected onto a translucent screen through a mirror attached to the head coil, and responses were recorded via two buttons held in their left and right hands.

In different runs participants performed either a delayed number comparison task, or a mental arithmetic task. In the delayed number comparison task (Fig 1A), different numbers presented either in symbolic or non-symbolic formats were presented in random positions inside a white circular region subtending ~7 degrees of visual angle at the center of the screen. Black Arabic digits and numbers of items were shown with two different fonts (Arial Rounded MT versus Times New Roman for symbolic numbers) and shapes (circles versus triangles for non-symbolic numbers). The total surface area covered (number of black pixels) was approximately equated between all non-symbolic numbers (resulting in smaller items for larger numerosities) and symbols. Examples of all conditions are shown in Figure 1B.

**Figure 1.**
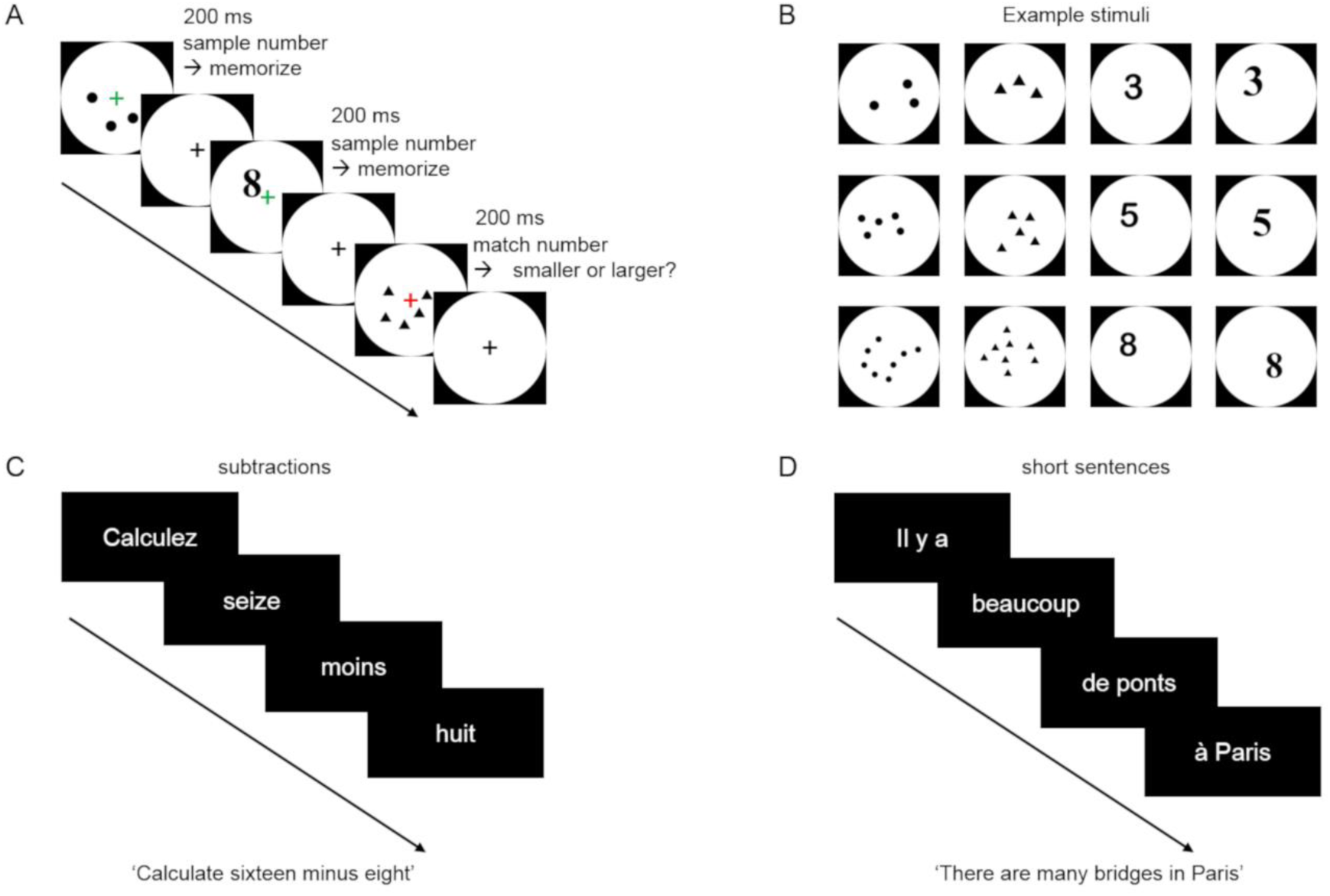
Experimental Paradigms. (A) In a delayed number comparison task, sample and comparison numbers were briefly shown (200 ms) in either non-symbolic or symbolic (Arabic digit) format. Participants were instructed to keep in memory the number seen in a given trial until the following trial appeared (after 10.5 s), and to perform a numerical comparison on occasional match trials, marked by a change in the fixation color. Participants were asked to judge whether the number displayed in the match trial was smaller or larger than the one seen in the previous sample trial. (B) Examples of sample stimuli. (C-D) During the mental arithmetic task, participants performed mental calculation (subtractions) according to written verbal instructions (C) or read math unrelated sentences (D).

The delayed comparison task started with brief (200 ms) presentation of a sample stimulus. Participants had to attend to the numerical content of each stimulus and to hold this information in memory until the following stimulus was presented (after an SOA of 10.5 s). One second before the onset of the following trial the fixation cross color changed from grey to either red or green. When red, this marked a match trial: in that case participants had to compare the current stimulus with the previously seen one, and respond by pressing one of the two buttons held by their left or right hand depending on whether they judged the current stimulus as numerically larger or smaller than the previous one. On the contrary, if the fixation cross turned green, participants only had to update their memory with the new sample stimulus. Three different sample numbers (digits 3, 5 or 8 or the corresponding numbers of items, see Fig. 1B for examples) were used and two possible match numbers could appear in each case (2 and 5 for sample 3, 3 and 8 for sample 5, 5 and 13 for sample 8). The presentation format (non-symbolic vs symbolic) always differed between a given sample and match. Each participant performed six 8.5 min long runs for the delayed number comparison paradigm. Each run contained six sample trials and two match trials (one smaller and one larger) per number and format.

In addition, all but one participant also performed a 4.9 min run with a mental arithmetic task (Fig 1C and D) adapted from a previously published functional localizer study (Pinel and Dehaene, 2010; Pinel et al., 2007). One participant was not tested with this paradigm, due to a longer than usual preparation procedure at the beginning of the session and subsequent lack of time. In different blocks, participants either solved mental subtraction problems according to verbal instruction (as for example: “Calculez quinze moins sept” [Calculate fifteen minus seven], see Fig 1C), with the first operand ranging from 10 to 19 and the second from 2 to 9, or read mathematics-unrelated sentences (as for example: “Il y a beaucoup de ponts à Paris” [There are many bridges in Paris], see Fig 1D). Each one of six blocks for each condition contained ten sentences, which were written in white on a black background, and centrally presented on four successive screens (each shown for 250 ms) separated by a 100 ms interval within sentence and a 2700 ms interval at the end of each sentence). Each screen presented a maximum of three words. Calculation and reading blocks were interleaved with baseline periods consisting of an additional 4 s of blank screen.

Stimuli were presented under Matlab 9.0 using Cogent (http://www.vislab.ucl.ac.uk/cogent.php) or using E-Prime software.

### Data Analysis

Statistical parametric mapping software (SPM12, https://www.fil.ion.ucl.ac.uk/spm/software/spm12/) was used to motion-correct the EPI images and to co-register them to the first single-band reference image. EPI images were corrected for distortions in FSL (https://fsl.fmrib.ox.ac.uk/fsl/fslwiki/FSL) in two steps: first we estimated a set of field coefficients with the topup function from the single-band reference images of the two initial volumes acquired with opposite phase encoding directions, and then we applied these to all the EPI images with the apply_topup function. Freesurfer (https://surfer.nmr.mgh.harvard.edu/) was used to perform cortical surface reconstruction of the anatomical image and boundary based registration of the mean single-band reference image to each subject’s anatomy.

The preprocessed EPI images (in subjects’ native space) were entered into GLMs using SPM, for the delayed comparison task modelling separately 12 sample stimulus conditions (3 numbers × 2 formats × 2 stimulus sets [shapes/fonts in case of non-symbolic/symbolic format]) within each run and 4 match stimulus conditions (2 formats × 2 magnitudes [smaller vs larger than sample]) as stick functions (using the default of 0 duration for events) convolved with the standard hemodynamic response function. Only two regressors (calculation and reading) modelling the onset of each sentence with a duration of 3.5 s were included in the GLM for the mental arithmetic task. To account for serial auto-correlation, an AR(1) model was used and low-frequency signal drifts were removed by high-pass filtering the data with a cutoff of 128 s.

To identify the cortical areas preferentially involved in perception of non-symbolic numbers, we contrasted the activation elicited by non-symbolic against symbolic sample stimulus conditions during the delayed number comparison task (contrast name: ‘Non-symbolic > Symbolic’). To isolate the correlates of two different numerical operations (comparison and calculation), we contrasted (A) the activation elicited by all match stimulus conditions against all sample stimulus conditions (contrast name: ‘Comparing > Viewing’), and (B) the activation elicited while participants performed mental subtractions against the activation elicited while reading mathematical unrelated sentences (contrast name: ‘Calculation > Reading’). These three contrasts were first created in each single subject’s volume space and then projected onto the surface with Freesurfer. Single subject’s contrast maps were aligned to fsaverage and smoothed with a 3-mm fwhm Gaussian kernel. Finally a random-effects group analysis was performed in the surface space. The resulting statistical maps were thresholded at p<.05 corrected for multiple comparisons at cluster level with cluster forming threshold p<.001.

Exploratory individual surface-based statistical analyses were also performed for visualization purposes, by using the design matrices previously configured in SPM, and estimating the GLM on each subject’s cortical surface using Freesurfer’s mri_glmfit function. For each subject and contrast we then selected the 2% most significant vertices, binarized their value and assigned them a color depending on the contrast (red for the voxels mostly activated in the contrast ‘Non-symbolic > Symbolic’, blue for ‘Comparing > Viewing’ and green for ‘Calculation > Reading’). Voxels activated in more than one contrast were marked by intermediate color (yellow for ‘Calculation > Reading and Non-symbolic > Symbolic’, cyan for ‘Calculation > Reading and Comparing > Viewing).

To statistically compare the contributions of the different contrasts across sub-regions of parietal cortex, we further performed an across-subjects regions-of-interest (ROI) analysis on individual subject’s statistical maps. For each subject, we defined anatomical regions of interest from two surface based parcellation schemes: one based on the Destrieux et al. (2010) atlas, which identifies the major sulci and gyri based on curvature estimates (Fig. 2A), and the other based on the Wang et al. (2015) atlas, which provides probabilistic maps of visual field map representations, including those from IPS0 to IPS5 (Fig. 2B). All ROIs were created on the Freesurfer surface and projected back into each subject’s volume space.

**Figure 2.**
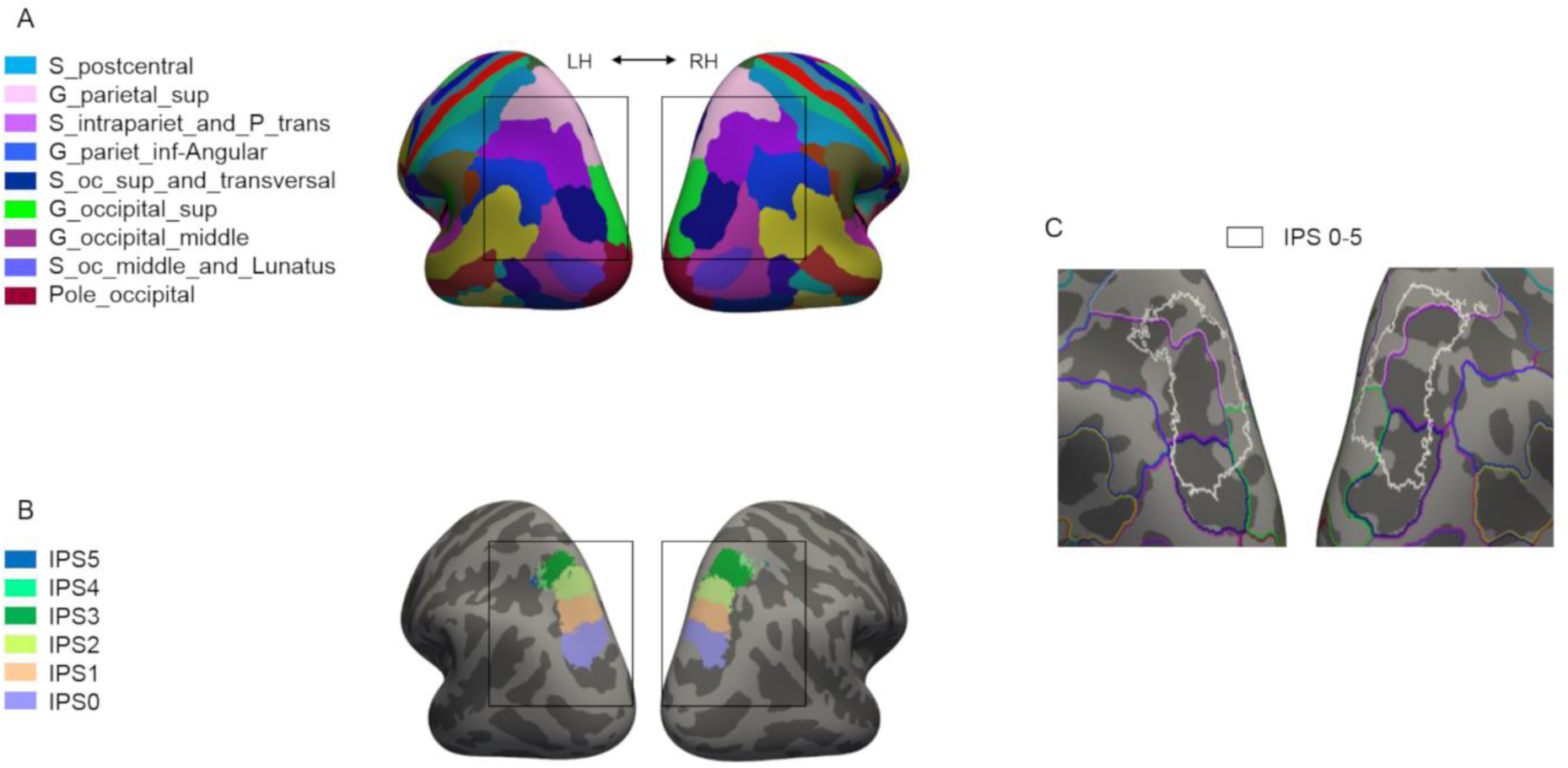
Visualization of regions of interest. ROIs based on (A) Freesurfer anatomical parcellation according to Destrieux et al. (2010) atlas and (B) visual field map representations IPS0 to IPS5 derived from the Wang et al. (2015) atlas are shown color-coded on the inflated template brain. The brain regions enclosed within the black rectangle are shown in more detail in (C) where ROIs defined by the two atlases in (A) and (B) are superimposed. The representation of visual field map ROIs IPS0 to IPS5 (IPS0-5 complex, white outline) overlaps with the superior parietal gyrus ROI (SPG, pink outline), the intraparietal sulcus and transverse parietal sulci ROI (IPS, purple outline), the superior occipital sulcus and transverse occipital sulcus ROI (blue outline) and the superior occipital gyrus ROI (green outline).

We focused the ROI analysis on parietal cortex (enclosed within the black rectangles in Fig. 2A and B), merging the left and right hemispheres. The parietal field map representations IPS0 to IPS5 derived from the Wang et al. (2015) atlas were merged into one large ROI (IPS 0-5 complex). As highlighted in Fig 2C, these visual field maps (IPS 0-5 complex, white outline) partly overlap with several gyri and sulci of the parietal cortex, including the Destrieux Atlas intraparietal and transverse parietal sulci (IPS) and the superior parietal gyrus (SPG), without fully matching any of them. We subdivided the region surrounding the fundus of the IPS into four ROIs roughly extending from lateral to medial, or inferior to superior parietal lobule: 1) Destrieux Atlas intraparietal sulcus exclusive of IPS0-5, 2) Destrieux Atlas intraparietal sulcus inclusive of IPS0-5, 3) Destrieux Atlas Superior Parietal Gyrus inclusive of IPS0-5, and 4) Destrieux Atlas Superior Parietal Gyrus exclusive of IPS0-5. For each subject, mean t-scores for the different contrasts were extracted from these four ROIs, as well as from further ROIs comprising the entire IPS0-5 complex, or its separate subparts: IPS0, IPS12 (merging IPS 1 and 2) and IPS345 (merging IPS 3, 4 and 5). As a measure of regionally specific contributions more independent of differences in overall activation strength across different contrasts, for each ROI and contrast, we also computed the difference between the mean t-scores measured inside and outside each ROI (i.e. in the rest of the parietal lobe, here defined by the union of the following Destrieux Atlas regions: Superior Parietal Gyrus, Angular Part of Inferior Parietal Gyrus, Supramarginal Part of Inferior Parietal Gyrus, Postcentral Sulcus, and Intraparietal Sulcus). Differences in signal strength across ROIs and contrasts were statistically tested with repeated measures ANOVAs.

## Results

To identify brain regions preferentially recruited during different types of numerical processing, such as numerosity perception, numerical comparison, and calculation, as a first step, we performed surface-based group analyses. Fig 3 shows the main three different contrast maps displayed on the surface of a template brain in relation to the parcellations derived from the two atlases used.

**Figure 3.**
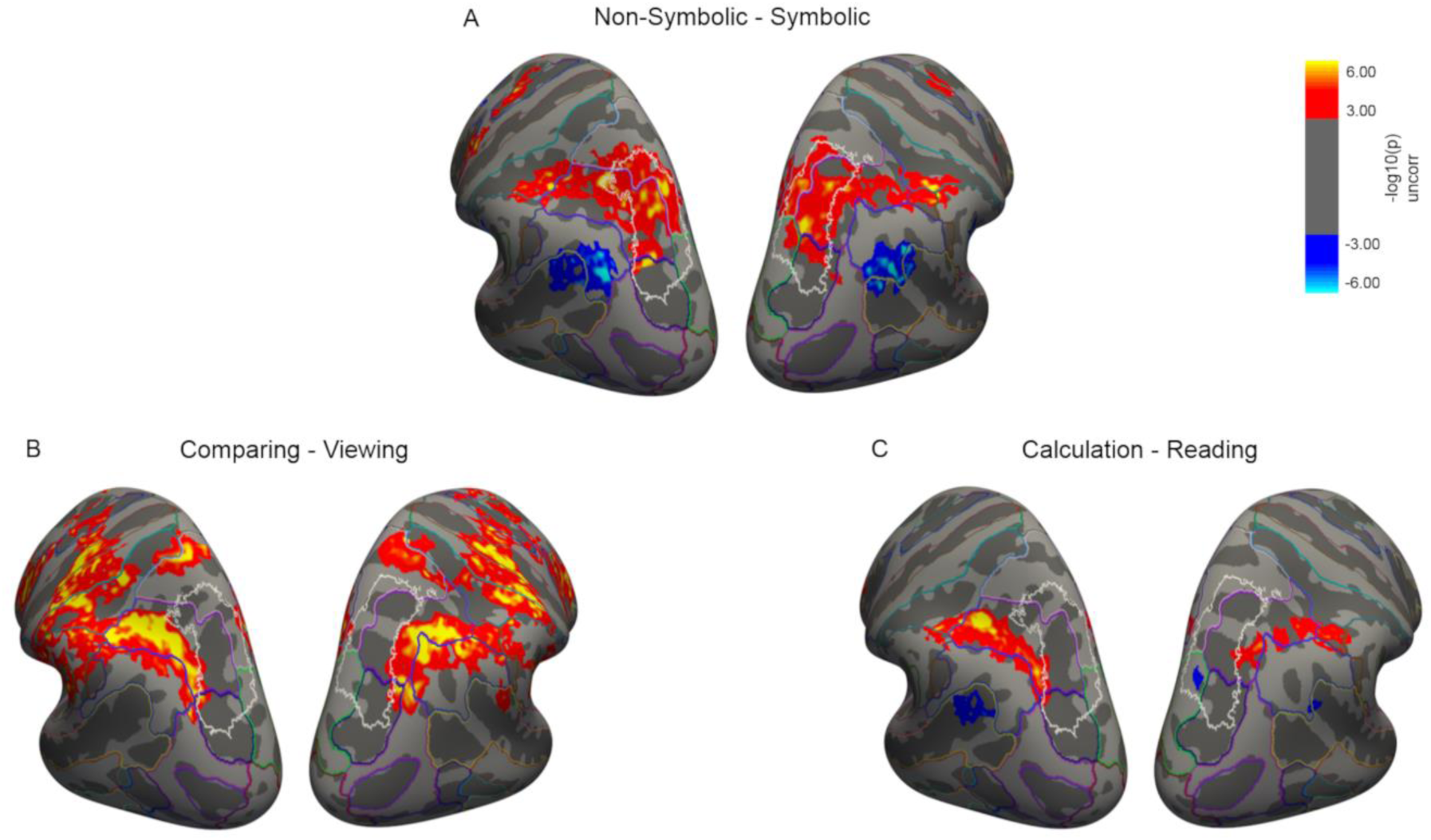
Regions within the intraparietal sulcus recruited for numerosity perception, numerical comparison and calculation – group maps. Activation maps from the surface-based random effects group analyses (n=15), thresholded at p<.05 corrected for multiple comparisons at cluster level with cluster forming threshold p<.001. Colour code corresponding to voxel-level significance (i.e. each voxel included in the clusters surviving correction is displayed with its uncorrected significance value). (A) Activations for mere viewing of “Non-symbolic > Symbolic” numbers occurred predominantly within the field map representations, while the reverse contrast showed activations in angular gyrus and superior temporal sulcus. (B) Activations for “Comparing > Viewing” of numbers were more pronounced in the areas outside the field map representations (in the intraparietal sulcus, inferiorly/laterally to the IPS0-5 complex). (C) Activations for “Calculation > Reading” were mainly found in regions outside the field map representations, while activations for the reverse contrast occurred in superior temporal sulcus.

Perception of non-symbolic numbers preferentially activated both occipital-parietal and frontal regions (red activations for the ‘Non-symbolic > Symbolic’ map in Fig 3A). More specifically, activations covered the superior occipital sulcus and transverse occipital sulcus, intraparietal sulcus and transverse parietal sulci (IPS), superior parietal gyrus (SPG), postcentral sulcus and precentral sulcus in the frontal cortex. Importantly, the parietal activations were mainly localized within the field map representations (delimited by the white outlines in Fig 3), covering the superior/medial portion of IPS and the inferior part of SPG. The reverse contrast showed that symbolic numbers elicited stronger activations than non-symbolic numbers (blue activations in Fig 3A) in the angular gyrus and superior temporal sulcus.

Explicitly performing a numerical comparison over mere viewing of sample numbers most strongly activated regions in the inferior/lateral bank of IPS, outside the visual field map representations (‘Comparing > Viewing’, Fig 3B). Activations for this contrast spread also more anteriorly into the postcentral sulcus and gyrus, and the central and precentral sulci. Mental calculation over reading also activated inferior/lateral regions of IPS (red activations for the ‘Calculation > Reading’ map in Fig 3C), while the reverse contrast led to some minor activations in the superior temporal sulcus (blue activations in Fig 3C).

Overall, the surface-based group analyses revealed that while all the different contrasts targeting different components of numerical processing activated areas within and around the IPS, different sub-regions within this larger area were activated predominantly as a function of the contrast: the medial/superior portion of the sulcus were most strongly recruited for mere perception of non-symbolic over symbolic numbers, whereas the most lateral/inferior regions of the sulcus were most strongly activated for numerical operations, i.e. during numerical comparison or calculation.

To further explore in how far the spatial organization of regional activations observed in the group analyses within sub-regions of intraparietal cortex was also evident at the level of individual subjects, we visualized individual subjects’ activations on their corresponding cortical surfaces. Fig 4 shows four representative individual subjects’ maps where the most significant vertices for each contrast are shown together on the participants’ inflated surfaces. The topological organization of activations in the parietal cortex observed in the group analysis was evident also in individual subjects: the medial/superior sub-regions of IPS and the inferior portion of SPG, comprising the visual field representation, were activated during simple perception of non-symbolic over symbolic numbers (see red activations) while comparing numbers or performing mental calculation elicited activations within progressively more lateral/inferior sub-regions (blue and green activations respectively). Vertices activated by more than one contrast were located in intermediate regions.

**Figure 4.**
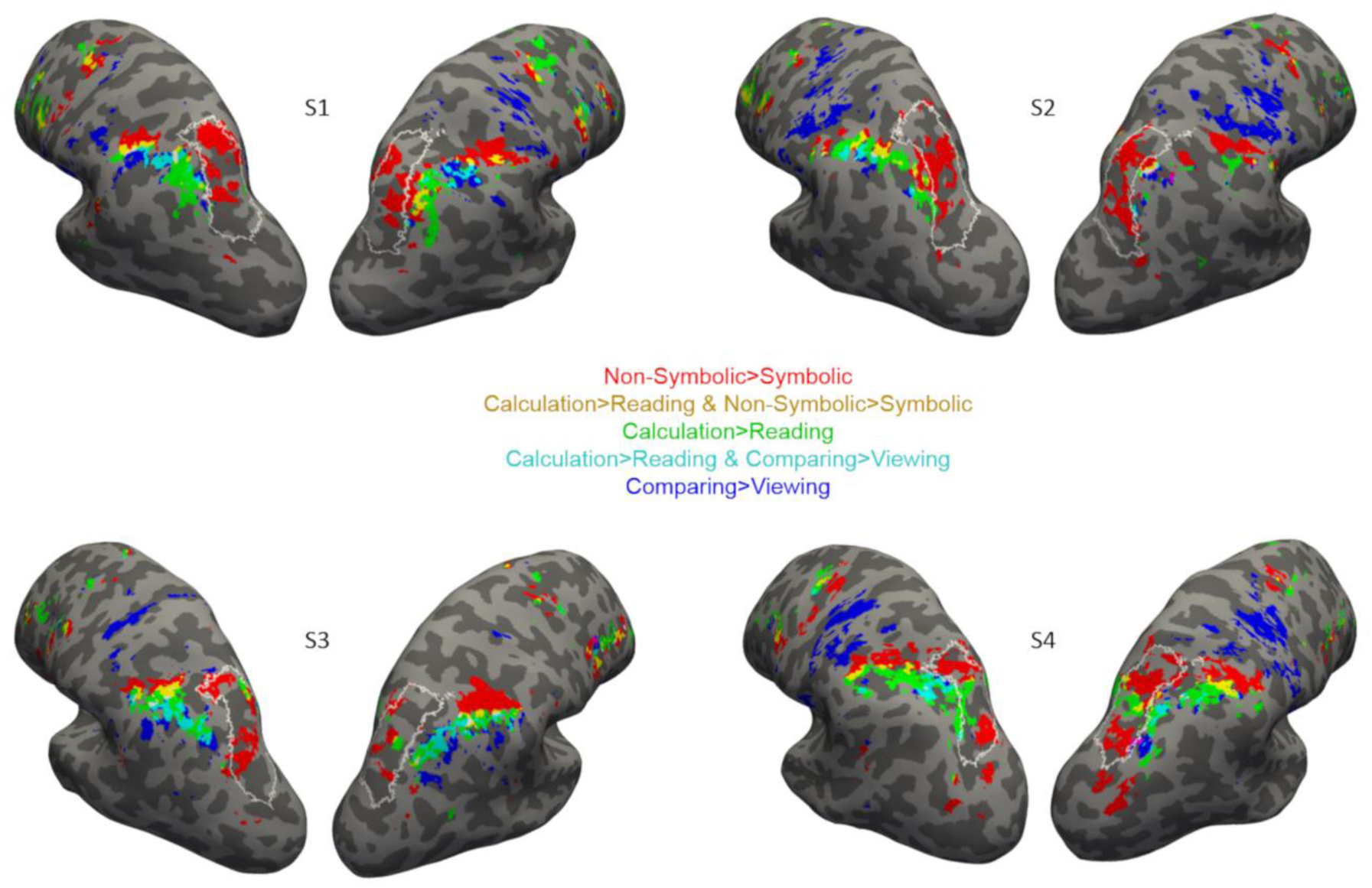
Regions within the intraparietal sulcus recruited for numerosity perception, numerical comparison and calculation - individual subjects’ maps. For each subject, maps display the 2% most significant vertices for each contrast (specified by colors, see legend) shown on the participants’ inflated surface. These representative individual subject maps indicated a similar topographical progression to the one observed in the group analyses. The white outline represents IPS 0-5 borders.

To statistically test at the level of individual subjects for systematic differences in sub-regions recruited as a function of the contrast, we performed additional ROI analyses of which the results are shown in Fig 5. First, we focused these ROI analyses specifically on parts of anatomically defined IPS and SPG that either did or did not overlap with the visual field map representations (Fig 5A). The IPS0-5 complex centrally overlapped with parts of both IPS and SPG which further extended laterally/inferiorly and medially/superiorly from the IPS0-5 complex, respectively. We therefore extracted the signal for the different contrasts (mean t-values across voxels for each individual subject) from four ROIs defined along a lateral-medial gradient. Specifically, moving from the most inferior/lateral to the most medial/superior part of the intraparietal region, we defined the first and second ROIs along IPS, excluding or including the IPS0-5 complex, respectively, then in the third and fourth ROIs the IPS 0-5 complex was included or excluded, respectively, from the ROIs defined along SPG.

**Figure 5.**
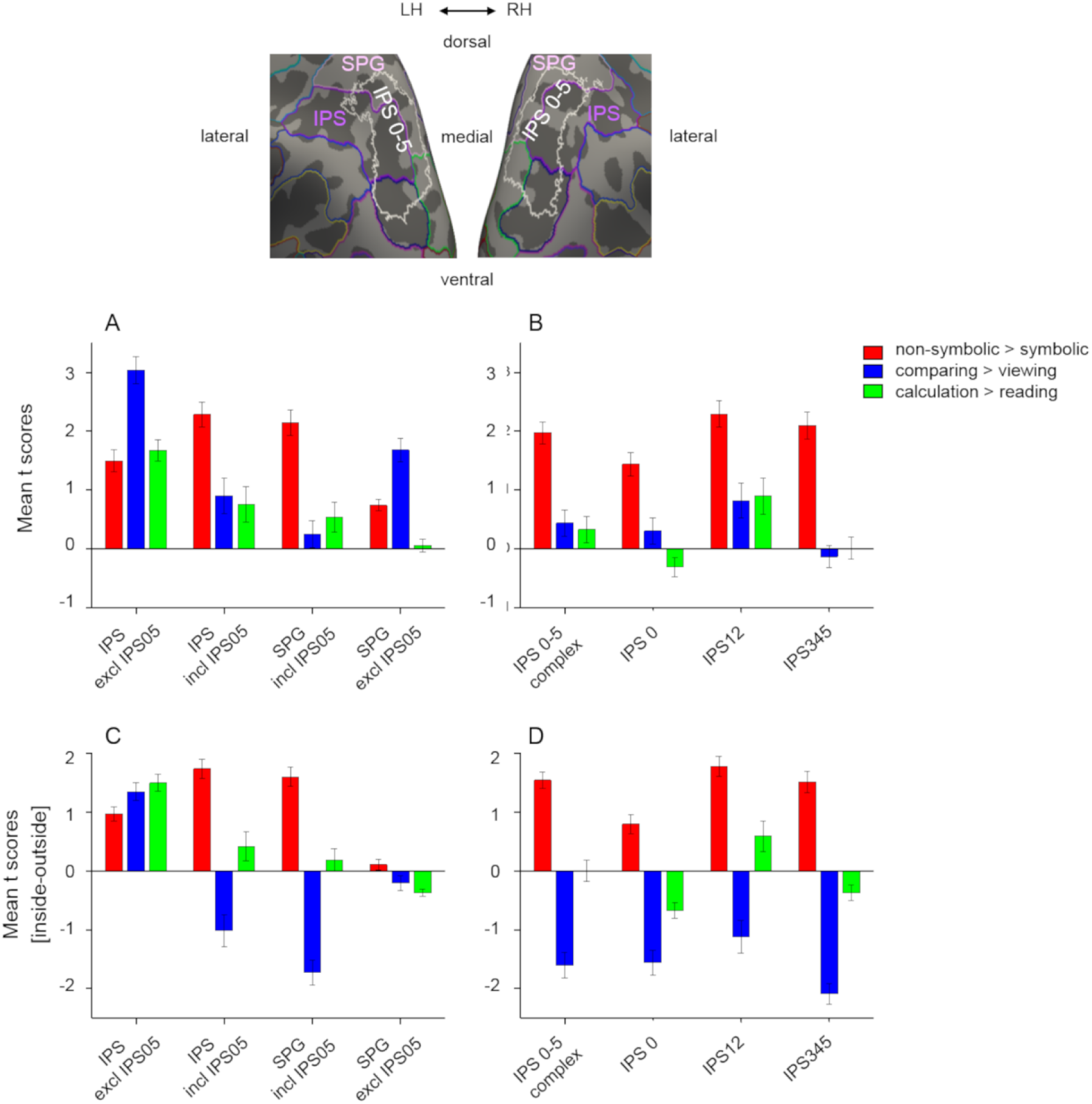
Results of ROI analyses. Mean t-scores for the selected ROIs corresponding to (A) anatomically defined intraparietal sulcus and superior parietal gyrus according to Destrieux et al. (2010) masked inclusively or exclusively with IPS0-5 fieldmap representations according to Wang et al. (2015) (see ROI display on top) and (B) visual field map representations IPS0-5 overall and its smaller subdivisions. (C and D) Differences in mean t-scores for the same ROIs with respect to the rest of parietal cortex (inside ROI-outside ROI). The ROI analyses confirmed a cross-over pattern: higher t-scores for “Non-symbolic > Symbolic” numbers (red bars) within IPS 0-5 field maps with respect to more lateral IPS areas, and higher t-scores for contrasts “Comparing > Viewing” (blue bars) and “Calculation > Reading” (green bars) in lateral IPS outside field maps with respect to IPS0-5 complex. Plots show mean t-scores (or difference of mean t-scores) across subjects (n=15) ± standard error of the mean (SEM).

Mean t-scores varied across the four ROIs depending on the contrast (Fig 5A), as confirmed by the significant interaction between ROI and contrast (F(3.6,50.6)=45.6,p<10^−5^). The mean t-scores for the contrast ‘Non-symbolic > Symbolic’ were significantly above zero in all four ROIs (all p<10^−5^), however the signal intensity varied between regions (Fig 5A, red bars): mean t-values became significantly higher when proceeding from the most lateral ROI in IPS, which did not include the IPS 0-5 complex, towards the more medial ROIs, which included the IPS 0-5 complex along IPS and SPG, respectively, and then significantly decreased again for the most medial ROI in SPG, in which the IPS 0-5 complex was excluded (post-hoc tests across ROIs: all significant at p<0.0005 at least, except for the difference between the two ROIs including the IPS 0-5 complex that was not statistically significant). On the contrary, the mean t-scores measured for the contrast ‘Comparing > Viewing’ followed an opposite trend, being highest whenever the IPS 0-5 complex was excluded from the ROI, i.e. for the most lateral ROI along IPS and the most medial ROI along SPG, and lowest for the IPS and SPG ROIs inclusive of the IPS0-5 complex (the latter not even being significantly different from zero, Fig 5A, blue bars, post-hoc tests across ROIs were all significant with p=0.01 at least). Finally the mean t-scores for the contrast ‘Calculation > Reading’ were highest in the most lateral ROI defined along IPS, excluding the IPS 0-5 complex, and progressively and significantly lower in the more medial ROIs, reaching a value not significantly different from zero for the most medial ROI in SPG, excluding IPS 0-5 complex (Fig 5A, green bars, post-hoc tests for the most lateral vs the progressively more medial ROIs: p=0.003, p=0.0002, p<10^−7^).

To detect any potential additional specializations within the visual field map representations, we also compared mean t-scores for smaller subdivisions of the IPS 0-5 complex (in particular IPS0, IPS12, and IPS 345) (Fig 5B). The ANOVA showed a significant interaction between ROI and contrast (F(2.3,32.7)=5.5 p=0.007). The values measured for the contrast ‘Non-symbolic > Symbolic’ were significantly above zero in all ROIs (all p<10^−5^), and significantly higher in IPS12 and IPS345 with respect to IPS0 (post-hoc tests across ROIs: significant at p=0.002 at least). The mean t-scores measured for the contrasts ‘Comparing > Viewing’ and ‘Calculation > Reading’ were not significantly different from zero, except for IPS12 where nevertheless the t-scores for these contrasts were much lower than the ones measured for the contrast ‘Non-symbolic > Symbolic’ (post-hoc tests across tasks in IPS12: p=0.002 and p=0.003).

As an alternative measure of relative regional preference more independent of the overall activation strength of each contrast, we further computed for each contrast the difference between the mean t-scores measured inside each ROI and outside it in the rest of parietal cortex (Fig 5 C and D). Positive values indicate stronger activations inside a given ROI with respect to the rest of the parietal cortex, whereas negative values point at the opposite pattern. A significant interaction between ROI and contrast (F(3.7,51.9)=43.7,p<10^−5^) was confirmed also for these measures for the four main regions of interest. For the contrast ‘Non-symbolic > Symbolic’, mean t-score differences were positive inside IPS, especially when the IPS 0-5 complex was included in the ROI (Fig 5C), indicating higher values within these ROIs compared to the rest of parietal cortex. As for the SPG ROIs, the mean t-score differences were positive only when including the IPS 0-5 complex, whereas excluding it resulted in values not significantly different from zero. Regional differences were confirmed by significant post-hoc tests across ROIs inclusive versus exclusive of the IPS 0-5 complex (all significant at p=0.001 at least). For the contrast ‘Comparing > Viewing’, mean t-scores differences were positive in the most lateral ROI defined along IPS excluding IPS 0-5 complex, indicating higher values within the ROI with respect to the rest of parietal cortex, whereas more medial ROIs along IPS and SPG showed progressively more negative values, indicating a predominant activation in parietal areas excluding the IPS 0-5 complex (post-hoc test across ROIs all significant at p=0.02 at least). In the most medial ROI along the SPG, excluding the IPS0-5 complex, values were nearly zero. The mean t-score differences for the contrast ‘Calculation > Reading’ were positive and highest for the most lateral ROI defined along IPS excluding the IPS 0-5 complex (post-hoc tests for the most lateral vs the progressively more medial ROIs: p=0.002, p<0.0001, p<10^−7^), and they were not significantly different from zero for the other more medial ROIs, except for the most medial one where the value was negative.

Finally, for the IPS0-5 sub-regions a significant interaction between ROI and contrast (F(2.3,32.3)=5.9, p=0.005) was also confirmed for the t-score difference measure. The mean t-score differences for the contrast ‘Non-symbolic > Symbolic’ were positive for all ROIs, indicating a stronger activation inside the ROIs than in the rest of parietal cortex, although the difference was smaller for IPS0 (post-hoc difference between IPS0 and the other ROIs: significant at least at p=0.001). On the contrary, mean t-score differences for the contrast ‘Comparing > Viewing’ were always negative, and activation thus stronger in the rest of parietal cortex excluding the field map representations. The mean t-score differences for the contrast ‘Calculation > Reading’ were not significantly different from zero when considering the whole IPS0-5 complex, had a significantly negative value both in IPS0 and IPS345 and a significantly positive value in IPS12. However, the contrast’s values for the latter ROI were significantly lower than those for the ‘Non-symbolic > Symbolic’ contrast (post-hoc across contrasts for IPS12: p=0.003).

In sum, the ROI analyses confirmed a significant cross-over pattern: higher overall activations for the contrast “Non-symbolic > Symbolic” within IPS 0-5 field maps compared to most lateral IPS or most medial SPG areas, as well as compared to all the rest of parietal cortex, and higher overall activations for the contrasts “Comparing > Viewing” and “Calculation > Reading” in lateral IPS parts outside field maps compared to inside the field maps, and compared to all the rest of the parietal cortex. Within the IPS 0-5 complex a weak, but significant, sub-regional specialization emerged too: activations in IPS0 were lower than those in IPS12 and IPS345 for the contrast “Non-symbolic > Symbolic” and IPS12 showed low but significant activations also for the contrasts “Comparing > Viewing” and “Calculation > Reading”.

## Discussion

In the current study we tested whether numerical perception and different numerical operations activate overlapping brain areas, as previously suggested by coarse-scale quantitative meta-analyses (Arsalidou and Taylor, 2011; Arsalidou et al., 2018) or whether a finer-scale pattern of sub-regional specialization within the parietal cortex can be revealed by the enhanced spatial resolution provided by ultra-high field fMRI combined with cortical surface-based analysis. To more precisely localize the observed activation foci and shed new light on their potential relation with known sub-regions of macaque cortex, we further related activations to anatomical and functional markers on the cortical surface, with the help of two atlases based on curvature and visual topography.

Our results showed that numerical perception (mere viewing of non-symbolic numbers) and numerical operations (explicit comparison and calculation), all led to activations within and around the IPS, however with clear differences across conditions and sub-regions. Perception of non-symbolic numbers activated the superior/medial parts of IPS and SPG more strongly than symbolic numbers which in turn activated more the angular gyrus and superior temporal sulcus. On the other hand, operating on the numerical information either to perform a comparison task or to compute the result of simple subtraction problems maximally recruited different and more inferior/lateral areas of IPS with respect to those involved in numerosity perception.

Using population receptive field (pRF) mapping, Harvey et al. (2013) described a topographically organized map of preferential responses to non-symbolic numerosities that overlapped with the general area containing visual field map representations, even though not coinciding with the borders of any particular one of those maps. The specific paradigm with extensive amount of stimulation and long scanning time required to perform pRF mapping reliably made it unfeasible for us to use the same approach here. Rather, we used a more classical activation contrast for non-symbolic numbers (compared to symbolic numbers) overall. This contrast is similar to the ones used by some previous studies which also reported preferential activation for non-symbolic over symbolic numbers in intraparietal regions, either during explicit comparison (Holloway et al., 2010) or during mere viewing and memorizing of sample numbers (He et al., 2014), the latter case very comparable to the situation in our study. However, due to the relatively low spatial resolution used in those previous studies in combination with averaging across subject in whole brain space, they were not able to attribute these effects to specific subparts of the IPS, and we extend their results by localizing the effects more precisely to the superior/medial bank of the sulcus, to a large degree in overlap with visual field maps representations. In that sense, the results obtained with our contrast point into the same direction as the ones obtained by Harvey et al. (2013): regions responsive to perception of non-symbolic numbers on the upper bank of the IPS overlap with those showing visual topography, even though our contrast is likely to have recruited a somewhat wider set of regions than Harvey’s topographic numerosity maps per se.

In the current study, we did not limit our analysis to the activity elicited by numerosity perception, but we also compared the locations of this activation with the one elicited when a numerical comparison was performed as opposed to mere viewing of sample numbers. Congruently with previous studies using delayed comparison tasks (Cavdaroglu and Knops, 2018; Cavdaroglu et al., 2015) we observed that match trials, during which a comparison was made, elicited stronger signals in intraparietal areas with respect to sample trials. Once again, our analysis methods allowed us to more precisely pin down the location of these activations within the IPS than done previously, and compare it to the ones observed during mere perception. Comparison over mere viewing of numbers did not merely recruit the same regions to a stronger degree, but higher activity during comparison was found most strongly outside the IPS0-5 complex, and this part of the IPS located inferiorly and laterally to the field map representations was most strongly activated with respect to the rest of the parietal cortex.

The same inferior/lateral region of IPS was also more strongly activated by calculation over reading. It is important to note that only subtractions were tested in the current experiment and results need not be entirely identical for other types of arithmetical operations. Indeed, there is evidence suggesting that the neuronal correlates of different arithmetical operations show some heterogeneity (Chochon et al., 1999; Lee, 2000; Dehaene et al., 2003; Zhou et al., 2007; De Smedt et al., 2011; Prado et al., 2011; Rosenberg-Lee et al., 2011; Prado et al., 2014, though see: Kawashima et al., 2004). Neuropsychological cases of double dissociations between the ability to solve multiplications and subtractions (reviewed in Dehaene et al., 2003) have led to the suggestion that multiplications may be typically solved by recalling the solution from rote verbal memory, whereas subtractions may require actual computation based on some sort of internal manipulation of numerical quantities on an internal number line, possibly similarly to the strategy employed to solve numerical comparisons (Dehaene et al., 2003). Neuroimaging studies on healthy subjects have reported stronger IPS activations for subtraction with respect to multiplication (Chochon et al., 1999; Lee, 2000; Prado et al., 2011) or whenever a procedural strategy is used as opposed to fact retrieval in which case the angular gyrus is more involved (Polspoel et al., 2017; Tschentscher and Hauk, 2014).

In line with the idea of subtraction and comparisons involving potentially similar internal manipulations of quantity, we provide evidence for an overlapping neural substrate supporting these two operations, localized in IPS in the most inferior/lateral part of the sulcus, outside the field map representations which seem on the contrary more involved in numerical perception. Of course, the fact that a given region is similarly activated during two different tasks (such as number comparison and calculation here) at the univariate level, does not necessarily imply recruitment of identical neuronal populations. Interestingly, one previous study using multi-voxel pattern analyses found a significant correlation across voxels between the strength of numerical distance effects measured during a number comparison task and responses for subtraction over multiplication in the IPS, suggesting that the activation overlap extends to an intermediate scale of neuronal responses (Prado et al., 2011). Future studies should perhaps use related multivariate techniques to probe the neuronal codes underlying internally computed quantities, such as those representing the outcome of a comparison process or an arithmetical operation.

The main functional landmark in relation to which we mapped numerical processing related activity here are intraparietal field map representation which, as noted in the introduction, are considered the likely human equivalents of the macaque LIP/VIP complex within which neurons responsive to the numerosity of non-symbolic arrays have been described by neurophysiological studies (Nieder et al., 2006; Roitman et al., 2007). Identifying equivalence between areas is non-trivial related to the fact that human parietal cortex has differentially expanded and is also recruited by higher-level functions that are not present in monkeys, such as language, sophisticated tool use and higher-level mathematics (Grefkes and Fink, 2005; Kastner et al., 2017). Therefore, the number of areas and their relative localization with respect to IPS anatomy can show some differences across the two species, and suggestions for correspondence should rather emphasize similarities in characteristic functional response properties across areas. Such tentative functional equivalence has been proposed between lower-level intraparietal field map representations and LIP, as well as higher-level field map representations and VIP (Kastner et al., 2017; Konen and Kastner, 2008). In particular neurons in macaque LIP respond to saccadic eye movements (Andersen et al., 1990), whereas the majority of the neurons in VIP prefer smooth pursuit eye-movements (Schlack et al., 2003) and multisensory motion (Avillac et al., 2005). Similarly, in humans the responses across field map sub-regions change from IPS1/2, located in the posterior/medial parietal cortex, preferring saccadic eye movements, to IPS 3/4/5 preferring smooth pursuit eye movements (Konen and Kastner, 2008), located more anteriorly and laterally and roughly overlapping with areas responsive to visuo-tactile stimulation (Bremmer et al., 2001; Sereno and Huang, 2006), arguably most similar to macaque VIP.

The mentioned findings led us to investigate separately IPS0, IPS12 and IPS345 subparts here, and to consider our IPS12 and IPS345 ROIs as more likely corresponding to macaque LIP and VIP, respectively. However, we also note that this particular subdivision should be taken with some caution, as no one-to-one correspondence between individual regions in the two species may exist. So far, a higher number of field maps has been described in humans than in monkeys, and determining the exact equivalence between regions should take into account multiple criteria and is still a topic of ongoing research (Kastner et al., 2017). In the current study, the field map ROIs IPS12 and IPS345 both showed higher activations with respect to IPS0 and to the rest of parietal cortex during perception of non-symbolic numbers, in line with the preferential neuronal responses to non-symbolic numbers that have been described in macaque LIP and VIP.

In addition, significant, although less strong, activations where also observed in the IPS12 ROI during numerical comparison and calculation. Some responsiveness of superior parietal regions during numerical operations has been noticed previously (e.g. Dehaene et al., 2003) and been interpreted as reflecting attentional shifts along an imaginary number line. In line with this interpretation, activity in intraparietal regions identified by their responsiveness to saccadic eye movements could be read out to train a decoder to distinguish leftward from rightward saccades, and this decoder could subsequently be used to predict two different arithmetic operations (subtraction vs addition) presumably associated with leftward as opposed to rightward shifts along the number line (Knops et al., 2009). The fact that in our context we found operation-related activity within the IPS0-5 complex to be localized to IPS12, the most likely equivalent of macaque LIP which is preferentially involved in spatial attention and saccadic eye movements, fits well into this overall picture. However, beyond the minor result in IPS12, the regions most strongly recruited during numerical operations (both comparison and calculation) fell outside the field map representations, and into more lateral/inferior portions of the IPS. A correspondence between the areas maximally recruited during these types of operations and macaque regions VIP and LIP, as commonly assumed in the literature, therefore appears unlikely given the present results. Currently, it remains to some extent unclear which, if any, would be the counterpart of these more lateral human IPS regions in the macaque monkey brain. Interestingly, a functional connectivity study suggested the existence of evolutionarily novel cortical networks in humans for which no correspondence in the monkeys’ brain could be identified (Mantini et al., 2013). One of these networks which was in addition located within the areas having undergone the largest degree of cortical surface expansion between monkeys and humans, encompassed the intraparietal cortex near HIPS (Mantini et al., 2013), and could possibly overlap with the operation-related activations shown here on the inferior/lateral bank of the IPS.

In conclusion, intraparietal cortex is confirmed to play a crucial role in different components of numerical processing tasks, however, our study revealed a sub-regional specialization where more medial versus more lateral parts of the intraparietal sulcus are preferentially recruited during quantity perception (mere viewing of non-symbolic numbers) and numerical operations (comparison and calculation), respectively. A functional equivalence with the macaque LIP/VIP complex is likely for the former, but unlikely for the latter regions, located outside visual field map representations. In light of the current results it would be interesting to further investigate what is the more comprehensive functional response profile of the potentially human-specific lateral intraparietal sulcus subparts, and what might be the common computational denominator underlying different tasks recruiting these regions. Finally, future studies should also test whether the sub-regional specialization observed here in adults is already present in children or whether this differentiation emerges during development and mathematical learning.

## Acknowledgments

This work was supported by the French National Research Agency (grant number ANR-14-CE13- 0020-01 to E. E.).

We thank F. De Martino and V. Kemper for advice on fMRI acquisition parameters and procedures.

